# *Anopheles gambiae* mosGILT regulates innate immune genes and *zpg* expression

**DOI:** 10.1101/2023.08.01.551536

**Authors:** Gunjan Arora, Xiaotian Tang, Yingjun Cui, Jing Yang, Yu-Min Chuang, Jayadev Joshi, Andaleeb Sajid, Yuemei Dong, Peter Cresswell, George Dimopoulos, Erol Fikrig

## Abstract

Gene-edited mosquitoes lacking a <u>g</u>amma-interferon-inducible lysosomal thiol reductase-like protein, namely (*mosGILT^null^*) have lower *Plasmodium* infection, which is linked to impaired ovarian development and immune activation. The transcriptome of *mosGILT^null^ A. gambiae* was therefore compared to wild type (WT) by RNA-sequencing to delineate mosGILT-dependent pathways. Compared to WT mosquitoes, *mosGILT^null^ A. gambiae* demonstrated altered expression of genes related to oogenesis, 20-hydroxyecdysone synthesis, as well as immune-related genes. Serendipitously, the zero population growth gene, *zpg*, an essential regulator of germ cell development was found to be one of the most downregulated genes in *mosGILT^null^* mosquitoes. These results provide the crucial missing link between two previous studies on the role of *zpg* and *mosGILT* in ovarian development. This study further demonstrates that mosGILT has the potential to serve as a target for the biological control of mosquito vectors and to influence the *Plasmodium* life cycle within the vector.

## Introduction

Malaria remains one of the deadliest vector-borne diseases, with 247 million cases and approximately 619 thousand deaths reported globally in 2021 (Organization, 2022). It is caused by apicomplexan protozoa of the genus *Plasmodium* and transmitted by female *Anopheles* mosquitoes. During its journey through the vector, *Plasmodium* is profoundly influenced by mosquito factors that either facilitate or suppress parasite development (Arora et al., 2023; Dong et al., 2022; Shaw and Catteruccia, 2019; Waterhouse et al., 2007). In the mosquito midgut, *Plasmodium* development begins with the differentiation of gametocytes into gametes, followed by the formation of zygotes, ookinetes, and then oocysts. Parasite numbers suffer great losses, especially during traversal of the midgut epithelium (Baton and Ranford-Cartwright, 2005; Volohonsky et al., 2020). This midgut bottleneck of the *Plasmodium* life cycle is, therefore, an optimal target for blocking malaria transmission (Bennink et al., 2016; Blandin et al., 2008; Smith et al., 2014; Vijay et al., 2018; Wang and Jacobs-Lorena, 2013; Whitten et al., 2006). Considering the increase of insecticide resistance in *Anopheles* (Benelli and Beier, 2017; Hemingway et al., 2016; Ranson and Lissenden, 2016; Shaw and Catteruccia, 2019), a better knowledge of how mosquito-derived factors influence midgut-stage parasites is important for developing novel strategies for malaria intervention.

Blood-feeding is important for the female mosquito to obtain the nutrients required for egg development. Studies suggest that *Plasmodium* maturation in the midgut is influenced by concurrent mosquito reproductive processes, yet the underlying mechanisms remain poorly understood (Costa et al., 2018; Marcenac et al., 2020). Through CRISPR/Cas9 gene editing, our group demonstrated that disruption of a <u>mos</u>quito <u>g</u>amma-interferon-inducible lysosomal thiol reductase-like protein - *mosGILT* - caused underdeveloped ovaries with a reduction in egg development and the production of 20-hydroxyecdysone (20E), and vitellogenin (Vg) (Yang et al., 2020). Interestingly, *mosGILT*-deficient (*mosGILT^null^*) *Anopheles gambiae* also showed significant suppression of both *Plasmodium berghei* and *Plasmodium falciparum* infection, as measured by reduced oocyst intensity and prevalence (Yang et al., 2020). In parallel, another study showed that *Anopheles* mosquitos with a disrupted <u>z</u>ero-<u>p</u>opulation <u>g</u>rowth - *zpg* - gene also exhibited underdeveloped ovaries that produced low amounts of 20E and were less permissive to *P. falciparum* infection (Werling et al., 2019). Though both *mosGILT*^null^ and *zpg*^null^ mosquitoes exhibited somewhat similar phenotypes, it is unclear whether their mechanisms of action are similar or distinct. In this study, we elucidate the *mosGILT*-dependent global transcriptional responses in *A. gambiae*. The information here may provide novel insights into the *A. gambiae* reproductive system network, as well as innate responses associated with *Plasmodium* infection.

## Results

### Transcriptome analysis of *mosGILT^null^ A. gambiae*

To examine whether the molecular responses to a blood meal were altered in *mosGILT^null^ A. gambiae*, RNA sequencing (RNA-seq) was performed. *mosGILT^null^* and Wild type (WT, control) mosquitoes were fed on naïve mice and then separated into cardboard containers. The whole body, ovaries, and fat bodies were collected from blood-fed mosquitoes 24 h after completion of the meal. RNA-seq was carried out with total RNA extracted from the whole body, ovaries, or fat bodies from *A. gambiae* using Illumina sequencing technology. A total of 16 samples (from control and *mosGILT^null^ A. gambiae*) were sequenced (Supplementary Figure 1). After sequencing, each library generated around 10-28 million reads and approximately 78.0-99.7% of the reads in each library mapped to the *A. gambiae* genome (Supplementary Figures 2, 3, and 4).

Principal component analysis of mosquito libraries examined the clustering of data/samples after a blood meal. All biological replicates of control and *mosGILT^null^* samples were distributed into two distinct groups (Fig. 1A). Volcano plots indicated that there were differentially expressed genes (DEGs) in all three groups (Fig. 1B), with whole-body samples having the maximum number of DEGs (n=5749), followed by the ovaries and fat bodies. In the whole body, 108 DEGs were common to all three groups (Fig. 2A). Of particular interest were 4991 DEGs in the ovaries of *mosGILT^null^* mosquitoes which are potentially helpful to elucidate the role of mosGILT in *A. gambiae* development and *Plasmodium* infection.

**Figure 1:**
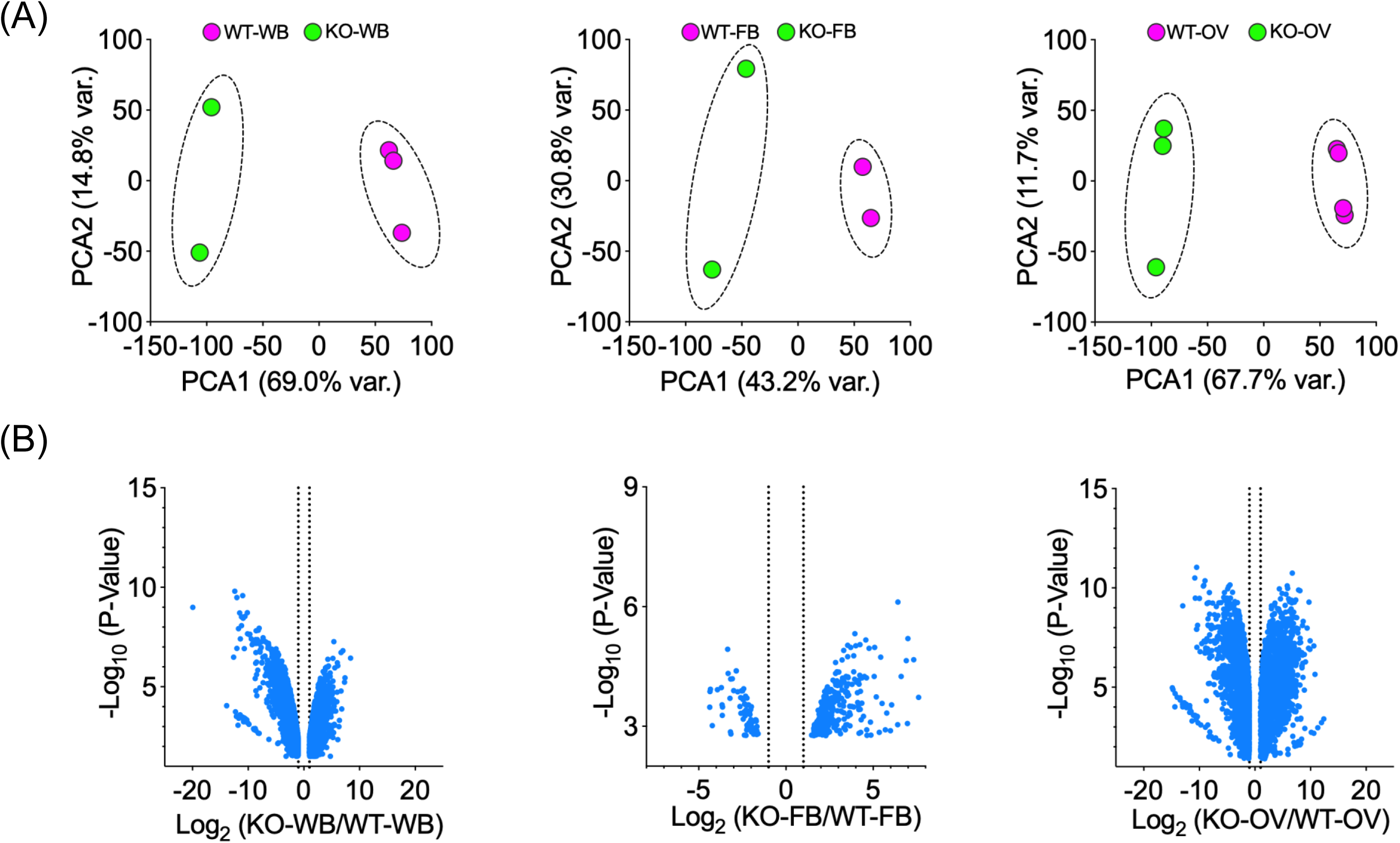
(A) Principal component analysis of the RNA-Seq data. The whole body (WB), fat body (FB) and ovary (OV) samples were collected at 24 h from blood-fed (BF) wild-type (WT) and mosGILT^null^ *Anopheles gambiae.* (B) Volcano plot analysis of differentially expressed genes (DEGs) of whole body (WB), fat body (FB) and ovary (OV) between wild type (control) and *mosGILT^null^ Anopheles gambiae*. DEGs with False Discovery Rate (FDR) *p*-values of < 0.05 and log_2_ fold changes > 1 are presented.

**Figure 2:**
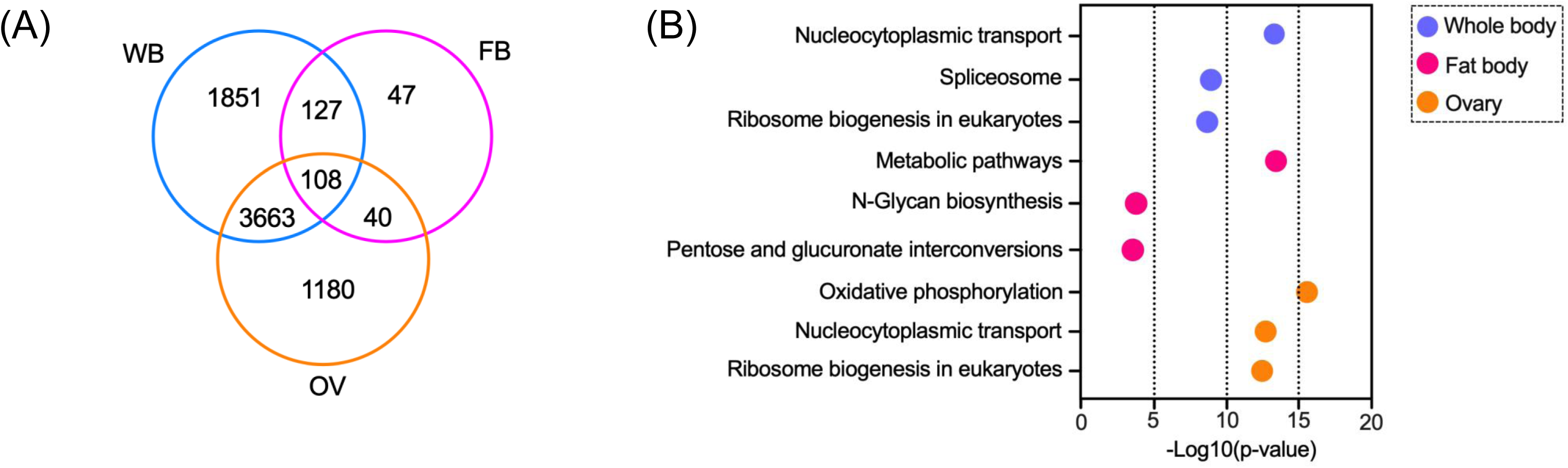
(A) Venn diagrams showing the number of differentially expressed genes (DEGs) between whole body (WB), fat body (FB) and ovary (OV) tissues of blood-fed *Anopheles gambiae*. The numbers in the overlapping areas indicate genes that were common to both or to all three different time points. The number of DEGs was highest between the whole body and ovary samples. (B) Bubble chart and circle graph showing enriched pathways in the whole body, fat body, and ovary of *mosGILT^null^* mosquitoes by KEGG (Kyoto Encyclopedia of Genes and Genomes) analysis. Gene ontology annotation and pathway enrichment analysis of the top differentially expressed genes (DEGs) in the whole body, fat body, and ovary are shown. Enriched functional ontology terms are shown on the left. The whole body, fat body, and ovary are represented by blue, red, and orange colors respectively. The Log_10_ (p-value) is shown on the X-axis.

### DEG analysis reveals that mosGILT has a role in nucleocytoplasmic transport

Since female *mosGILT^null^* mosquitoes display defects in ovarian development and show refractoriness to *Plasmodium*, we first focused on the nature of genes that showed differential expression patterns in the ovarian tissue. Among the 4991 DEGs, 117 genes had a more than a 100-fold increase in mRNA abundance compared to the WT controls, and 97 genes showed at least a 100-fold lower mRNA abundance. Several groups of genes showed altered expression patterns in oxidative phosphorylation and nucleocytoplasmic transport based on Gene Ontology (GO) annotation (Fig. 2B, and 3A, 3B). Nucleocytoplasmic transport (75 genes, p-value 2.60E-13) is a process crucial for cell function (Supplementary Table 1 and Table 1). Depletion of mammalian GILT (IFI30) in immune cells can inhibit cytokine production and the release of high mobility group box 1 (HMGB1), a nonhistone nuclear protein. The cytosolic translocation of the nuclear HMGB1 protein is then activated and associated with increased autophagy in GILT-depleted fibroblasts (Chen et al., 2022; Chiang and Maric, 2011). These findings suggest that mammalian GILT is involved in the homeostatic regulation of oxidative stress. In the absence of mosGILT, nucleocytoplasmic transport pathways are upregulated which can influence diverse cellular responses, such as autophagy. Indeed, autophagy negatively regulates Vg production in gonotrophic cycles in *Aedes* mosquitoes, which is consistent with the changes in Vg production shown by our group (Fig. 3C) (Bryant and Raikhel, 2011).

**Table 1.**

| ID | Source | Term ID | Term Name | P<sub>adj</sub> (query_1) |
| --- | --- | --- | --- | --- |
| 1 | GO:MF | GO:0003676 | nucleic acid binding | $3.175 \times 10^{-101}$ |
| 2 | GO:MF | GO:0005515 | protein binding | $5.581 \times 10^{-24}$ |
| 3 | GO:MF | GO:0015453 | oxidoreduction-driven active transmembrane t... | $4.107 \times 10^{-9}$ |
| 4 | GO:MF | GO:0015078 | proton transmembrane transporter activity | $3.330 \times 10^{-8}$ |
| 5 | GO:MF | GO:0003688 | DNA replication origin binding | $4.061 \times 10^{-5}$ |
| 6 | GO:MF | GO:0003712 | transcription coregulator activity | $6.201 \times 10^{-4}$ |
| 7 | GO:MF | GO:0140993 | histone modifying activity | $1.081 \times 10^{-3}$ |
| 8 | GO:MF | GO:0017056 | structural constituent of nuclear pore | $2.425 \times 10^{-3}$ |
| 9 | GO:MF | GO:0019887 | protein kinase regulator activity | $4.602 \times 10^{-3}$ |
| 10 | GO:MF | GO:0046933 | proton-transporting ATP synthase activity, rota... | $3.527 \times 10^{-2}$ |
| 11 | GO:BP | GO:0006139 | nucleobase-containing compound metabolic p... | $1.661 \times 10^{-94}$ |
| 12 | GO:BP | GO:0007049 | cell cycle | $1.334 \times 10^{-15}$ |
| 13 | GO:BP | GO:0009060 | aerobic respiration | $9.462 \times 10^{-14}$ |
| 14 | GO:BP | GO:0006913 | nucleocytoplasmic transport | $5.226 \times 10^{-12}$ |
| 15 | GO:BP | GO:0046034 | ATP metabolic process | $3.482 \times 10^{-4}$ |
| 16 | GO:BP | GO:0098660 | inorganic ion transmembrane transport | $1.117 \times 10^{-3}$ |
| 17 | GO:BP | GO:0070647 | protein modification by small protein conjugati... | $2.003 \times 10^{-3}$ |
| 18 | GO:BP | GO:0071824 | protein-DNA complex subunit organization | $4.031 \times 10^{-3}$ |
| 19 | GO:BP | GO:0098655 | monoatomic cation transmembrane transport | $4.079 \times 10^{-3}$ |
| 20 | GO:BP | GO:1902652 | secondary alcohol metabolic process | $6.149 \times 10^{-3}$ |
| 21 | GO:CC | GO:0005634 | nucleus | $2.120 \times 10^{-126}$ |
| 22 | GO:CC | GO:0070469 | respirasome | $9.751 \times 10^{-16}$ |
| 23 | GO:CC | GO:0016469 | proton-transporting two-sector ATPase complex | $3.651 \times 10^{-5}$ |
| 24 | GO:CC | GO:0005925 | focal adhesion | $3.592 \times 10^{-2}$ |

**Figure 3:**
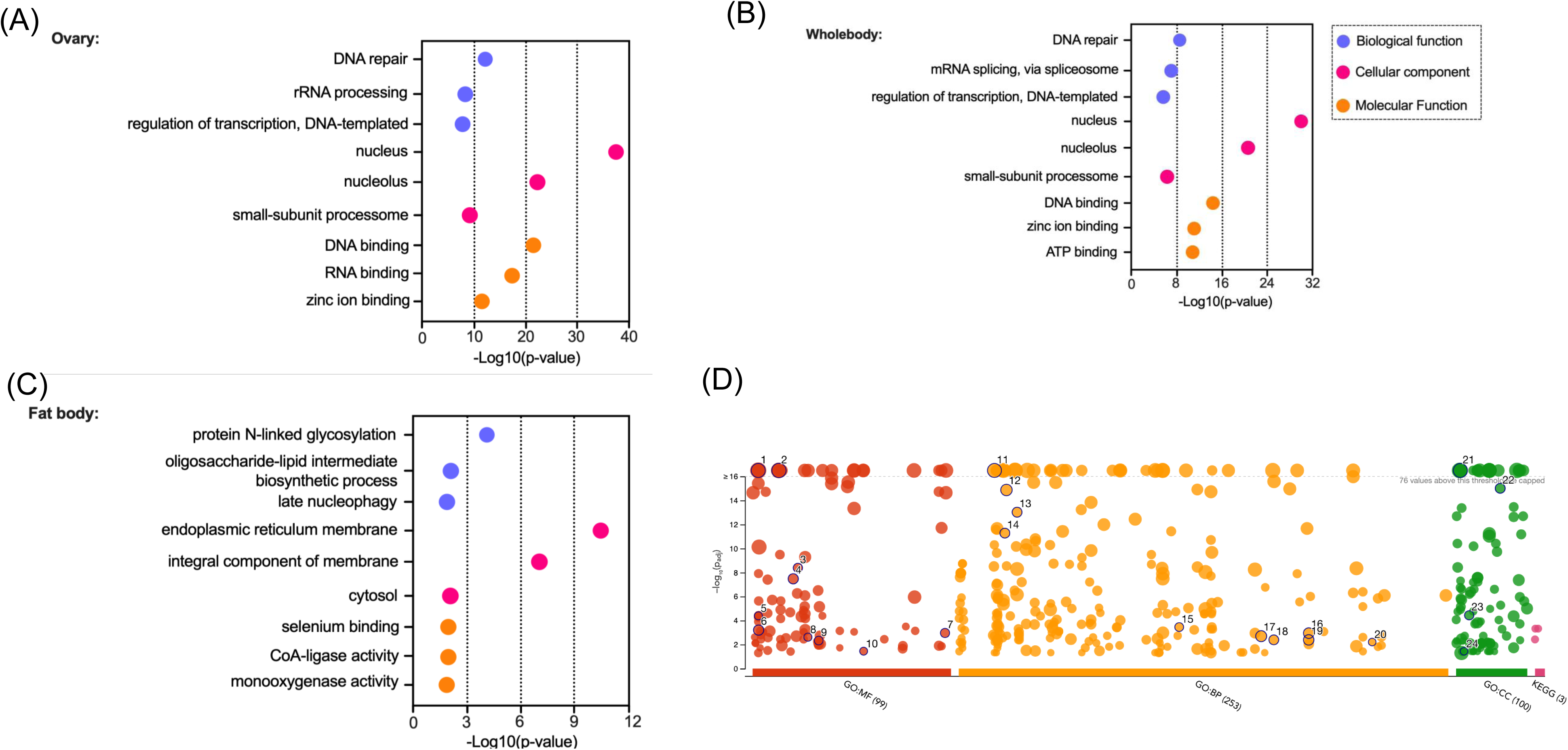
(A), (B) and (C) Bubble chart showing enrichment of Gene Ontology (GO) terms of the differentially expressed genes in the whole body, fat body, and ovary of *mosGILT^null^* mosquitoes. The biological processes, cellular components, and molecular functions are shown in blue, magenta and orange respectively. The –Log10 (p-value) for each process is shown on the X-axis. (D) The figure presents the Manhattan plot showcasing enrichment analysis of differentially expressed genes in the ovary of *mosGILT^null^* mosquitoes. The x-axis represents functional GO terms grouped and color-coded based on data sources. The y-axis displays adjusted enrichment p-values on a negative log10 scale.

### Whole body analysis of *A. gambiae* confirms the role of mosGILT in development and metabolism

Our previous studies demonstrated that mosGILT is implicated in embryonic and ovarian development and shows refractoriness to Plasmodium infection (Yang et al., 2020). To fully understand how female *mosGILT^null^* mosquitoes show resistance to *Plasmodium*, we also examined the genes that showed differential expression patterns in the whole body. Among the 6135 DEGs, 76 genes had a more than a 100-fold decrease in expression compared to the WT control, and 7 genes showed at least 100-fold upregulation. Similar to the ovaries, nucleocytoplasmic transport pathways were affected in the whole body of *mosGILT^null^* mosquitoes. The most downregulated proteins included AGAP007823 (meiosis arrest female protein 1, fold change: -1000000, p-value <0.0001) and AGAP008738 (eukaryotic translation initiation factor 4E transporter, fold change -14625, p-value 0.0001) which is involved in protein translation and oogenesis (Fig. 4A).

**Figure 4:**
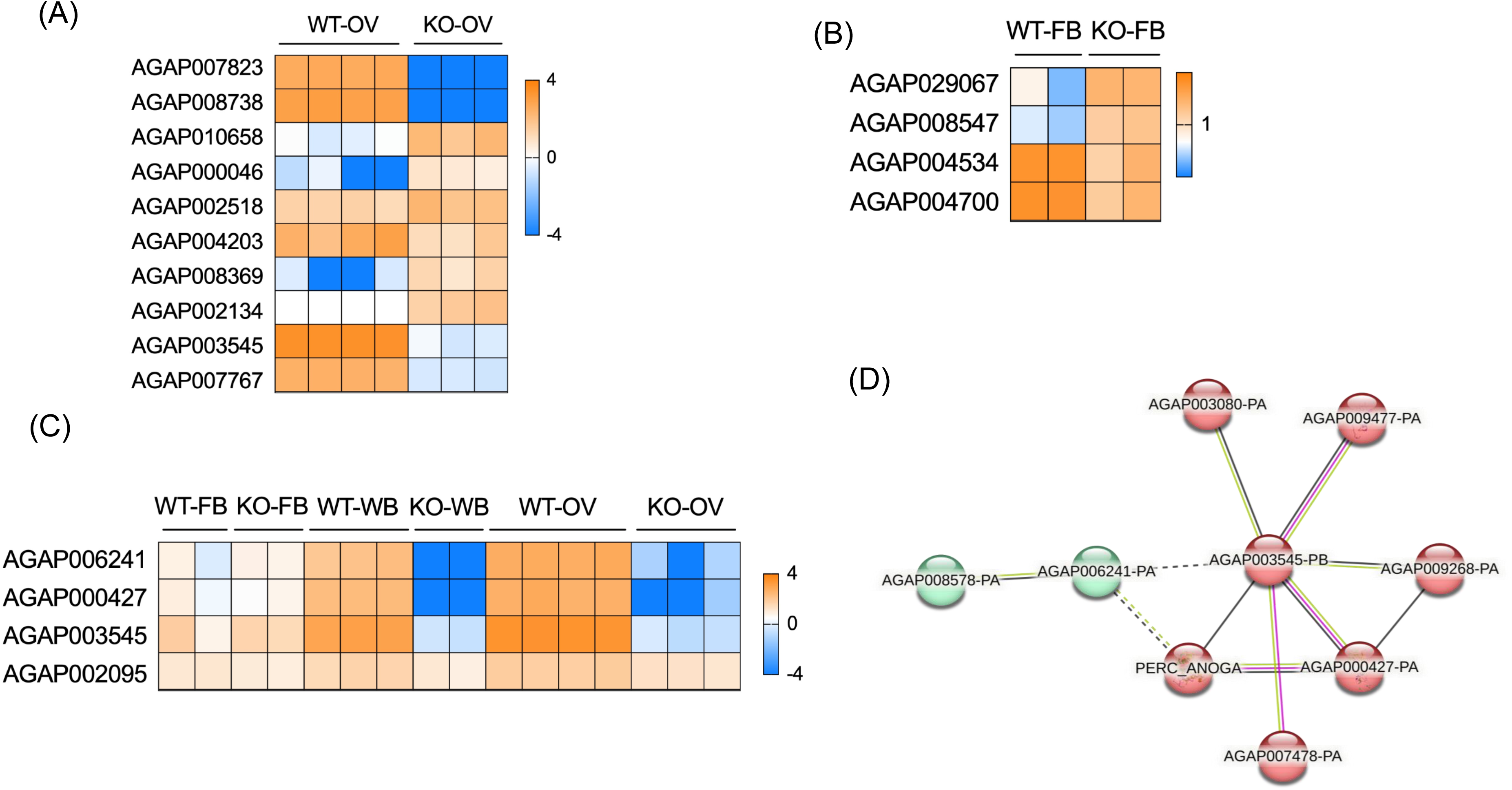
Heat Map analysis of key differentially expressed genes in the ovary (A), and fat body (B) of *mosGILT^null^* mosquitoes. (C) Heat map analysis of key differentially expressed genes that are related to germ cell development in *mosGILT^null^* mosquitoes. (D) STRING functional gene analysis of *zpg*-related genes: Each node represents a protein product of an investigated gene. Lines denote protein-protein interactions, with each edge representing an interaction including either physical or functional associations.

The most overexpressed genes included AGAP010658 (fold change 169, p-value < 0.00001) which encodes a homolog of the hexameric 2 beta protein or larval serum protein 1 which serves as a major storage protein. AGAP010658 supports *Plasmodium berghei* oocyst development and silencing of this gene by RNAi results in a significant increase in oocyst infection intensities and melanization of ookinetes (Lombardo and Christophides, 2016). Other highly expressed genes were AGAP000046 (a sugar transporter, fold change 160, p-value <00001) and AGAP002518 (delta l-pyrroline-5-carboxylate synthetase, fold change 102, p-value <.00001). While transcripts of AGAP004203 were decreased (fold change -343), another Vg-like protein was overexpressed (AGAP008369 fold change 102, p-value <0.00001). Among the major downregulated genes were AGAP002134 (vitelline membrane component which is required for the proper assembly of the eggshell) AGAP003545 (maternal effect protein oskar), and AGAP007767 (mosquito-specific bZIP1 transcription factor) (Fig. 4A). Indeed, silencing of *bZIP1* in the embryos of *Aedes aegypti* resulted in embryonic death (Criscione, 2013). These findings indicate that mosGILT is involved in both development and metabolism.

### mosGILT regulates metabolism in the *A. gambiae* fat body

Our previous study showed that *mosGILT* is expressed in the ovaries and fat body. In insects, the fat body plays central role in energy metabolism. After a blood meal, the fat body, which is the mosquito fat storage organ, initiates Vg synthesis. The fat body releases Vg into the hemolymph, and it is then taken up by developing oocytes via receptor-mediated endocytosis. Since *mosGILT^null^* mosquitoes showed lower Vg mRNA levels, we examined the transcriptional response in the fat body. Our analysis shows that in the fat body, gene transcripts of metabolic pathways (gene count: 93, p-value 6.30E-14) and pathways related to carbohydrate metabolism (pentose and glucuronate interconversions, fructose and mannose metabolism, galactose metabolism, biosynthesis of nucleotide sugars, glycolysis/gluconeogenesis and starch, and sucrose metabolism) demonstrated major changes in the *mosGILT^null^* mosquitoes. Among the most altered DEGs, AGAP029067 (lipid-binding protein) transcripts were enriched by a 194-fold while transcripts of AGAP008547, a gene with a predicted function in carbohydrate metabolism showed nearly 160-fold enrichment in the fat body of *mosGILT^null^* mosquitoes (Fig. 4A). Blood-meal ingestion therefore not only induces the expression of genes related to egg development but also increased genes related to oxidative stress.

In mammals, GILT is expressed in lysosomes and is required for optimal activity of cysteine proteases, such as cathepsin in macrophages (Balce et al., 2014; Ewanchuk et al., 2021). In mosquitoes, the fat body plays a central role in metabolism and nutrient storage, mirroring the functions of the liver in vertebrates. In response to blood-feeding, the *Aedes* fat body secretes a cathepsin-b-like thiol protease (Cho et al., 1999).

Interestingly, in our analysis of *mosGILT^null^ A. gambiae*, the most downregulated genes include proteases such as AGAP004534 (cathepsin B -like serine protease, fold change -18, p-value 0.0001), and AGAP004700 (trypsin-like serine protease, fold change -20, p-value 0.0003) (Fig. 4B).

### mosGILT regulates germ cell maintenance by controlling the expression of *zero population growth* (*zpg*)

*A. gambiae* females express *<u>z</u>ero <u>p</u>opulation <u>g</u>rowth* (*zpg,* AGAP006241), an essential gene for germ cell development. Genetic ablation of *zpg* in mosquitoes leads to severely underdeveloped ovaries and impaired egg production after a blood meal (Magnusson et al., 2011; Tazuke et al., 2002; Werling et al., 2019) similar to the *mosGILT^null^* mosquitoes. *zpg* and *mosGILT* are important for *Plasmodium* infection since mosquitoes lacking either gene show fewer oocysts after feeding on *P. falciparum*-infected blood compared to sibling controls. In the ovaries of *mosGILT^null^ A. gambiae*, *zpg* was strongly downregulated (fold change -8130, p-value 0.0001) which suggests *mosGILT* acts upstream of *zpg* or influences the molecular pathway that regulates *zpg* expression (Fig. 4C). We further performed STRING analysis (Fig. 4D) to determine whether mosGILT affected the expression of other genes that are associated with *zpg*. STRING analysis showed the genes that are in the *zpg* network and may be involved in germ cell development. Lack of mosGILT affected the expression of AGAP000427 (fold change -26814, p-value> 0.00001), which is in the *zpg* network and is a putative Vg receptor. In the ovaries of *mosGILT^null^*O mosquitoes, another *zpg*-related protein, *oskar* (AGAP003545) was strongly downregulated (fold change -8162, p-value> 0.00001) (Fig. 4C). Oskar is required for the formation of the germplasm in *Drosophila* (Ephrussi et al., 1991). Germplasm promotes the formation and specification of the primordial germ cells, the first cell lineage to form in the fertilized embryo (Kistler et al., 2018). Our genetic analyses of *mosGILT^null^* mosquitoes reveal underdeveloped ovaries and a decrease in 20E expression. Here we have analyzed genes involved in insect hormone biosynthesis and found that five genes (*AGAP001038, AGAP001039, AGAP004665, AGAP005992, AGAP000284*) involved in 20E biosynthesis were downregulated in *mosGILT^null^* mosquitoes (Fig. 5). Signaling of 20E is initiated by binding of 20E to the heterodimer ecdysone receptor (EcR)/ultraspiracle (USP) complex, which triggers a transcriptional response that promotes egg development. Our results show that *mosGILT^null^* also decreases the expression of ultraspiracle (USP, AGAP002095, fold change -4, p-value <0.00001) (Fig. 4C). The complex formed after 20E binding to EcR/USP upregulates the expression of an array of 20E-regulated genes which in turn controls the Vg transcriptional program (Ferracchiato et al., 2022). These results show that mosGILT regulates 20E levels, as well as the downstream transcriptional response.

**Figure 5:**
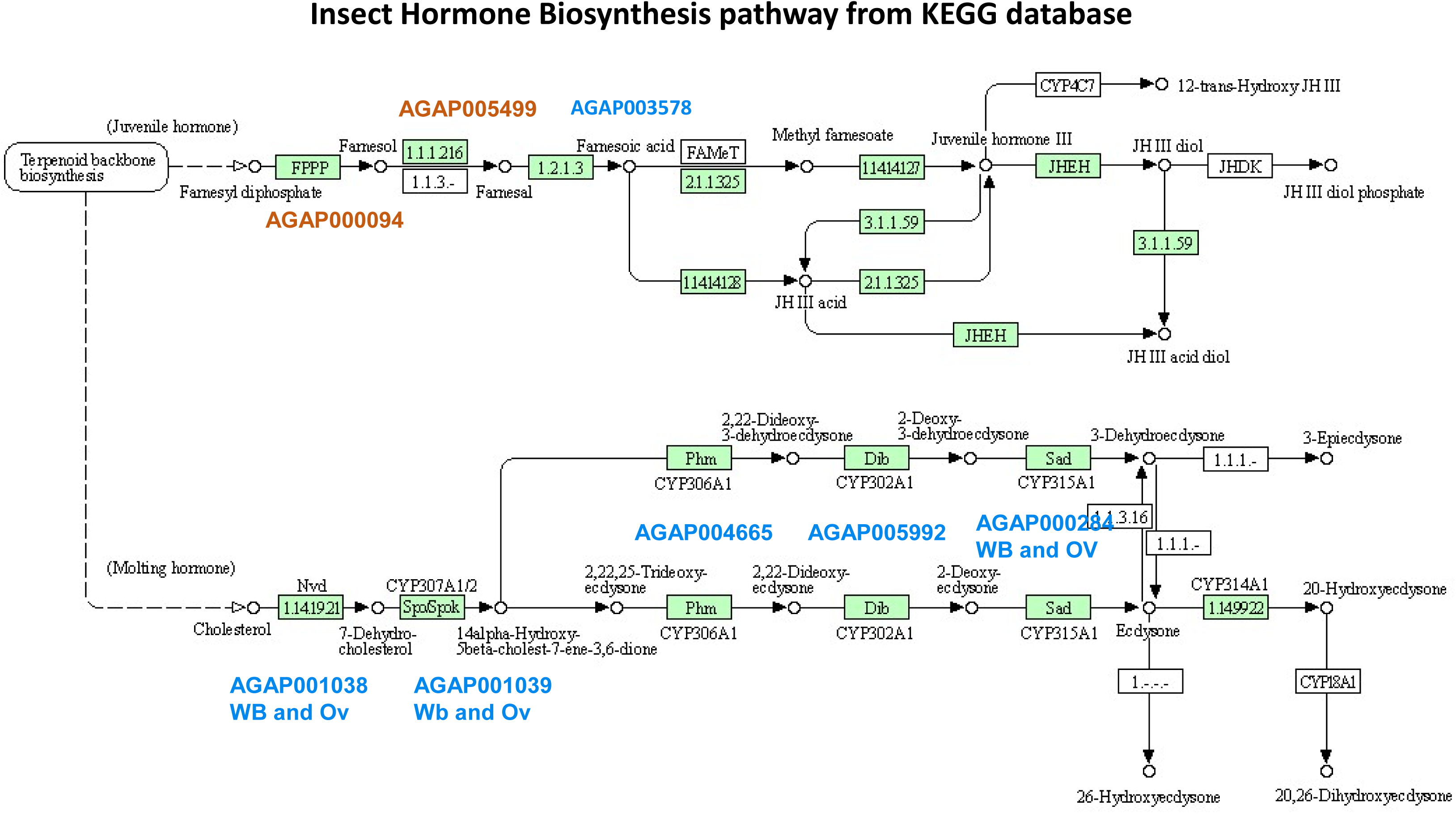
Insect hormone biosynthesis pathway in *Anopheles gambiae* based on KEGG pathway analysis. Enzymes are indicated in green highlight. The DEGs genes in *mosGILT^null^* mosquitoes are shown in orange (overexpression) and blue (downregulation) are indicated by the gene ids.

### Immune pathway alteration in *mosGILT^null^* mosquitoes

GILT was first identified as an IFNγ-inducible, lysosomal thiol reductase in mammals (Arunachalam et al., 1998). GILT is also key to antigen processing in mammals but in other organisms it is part of the innate immune response (Liu et al., 2013). Thioester-containing protein 1 (TEP1) is an important component of the *A. gambiae* innate immune system and plays a major role against *Plasmodium* parasites. TEP is an antimicrobial protein family that acts like the human complement protein C3 and can damage the cell membranes of pathogens. In our study, we observed an increase in TEP1 activity in *mosGILT^null^* mosquitoes (Yang et al., 2020). We also showed that strong refractoriness to *Plasmodium* infection in *mosGILT^null^* mosquitoes was related to increased TEP1 activity. Therefore, we examined immune genes that are differentially expressed in *mosGILT^null^* mosquitoes to better understand how TEP1 activity is influenced.

In our analysis, we found that AGAP028725 (SPCLIP1) was upregulated in the ovary (fold change 73, p-value 0.0001) and whole body (fold change 12, p-value 0.0001) of *mosGILT^null^* mosquitoes (Fig. 6). Clip domain serine proteases (CLIPs) play an important role in innate immunity in mosquitoes. These proteases are present in hemolymph and are involved in *Plasmodium* immunity, antimicrobial peptide synthesis and melanization. Previously it is shown that complement C3-like protein TEP1 mediated killing of *Plasmodium* is dependent on the CLIP-domain serine protease homolog SPCLIP1 (Povelones et al., 2013). SPCLIP1 is required for the accumulation of TEP1 on microbial surfaces, a reaction that leads to lysis of malaria parasites (Povelones et al., 2013).

**Figure 6:**
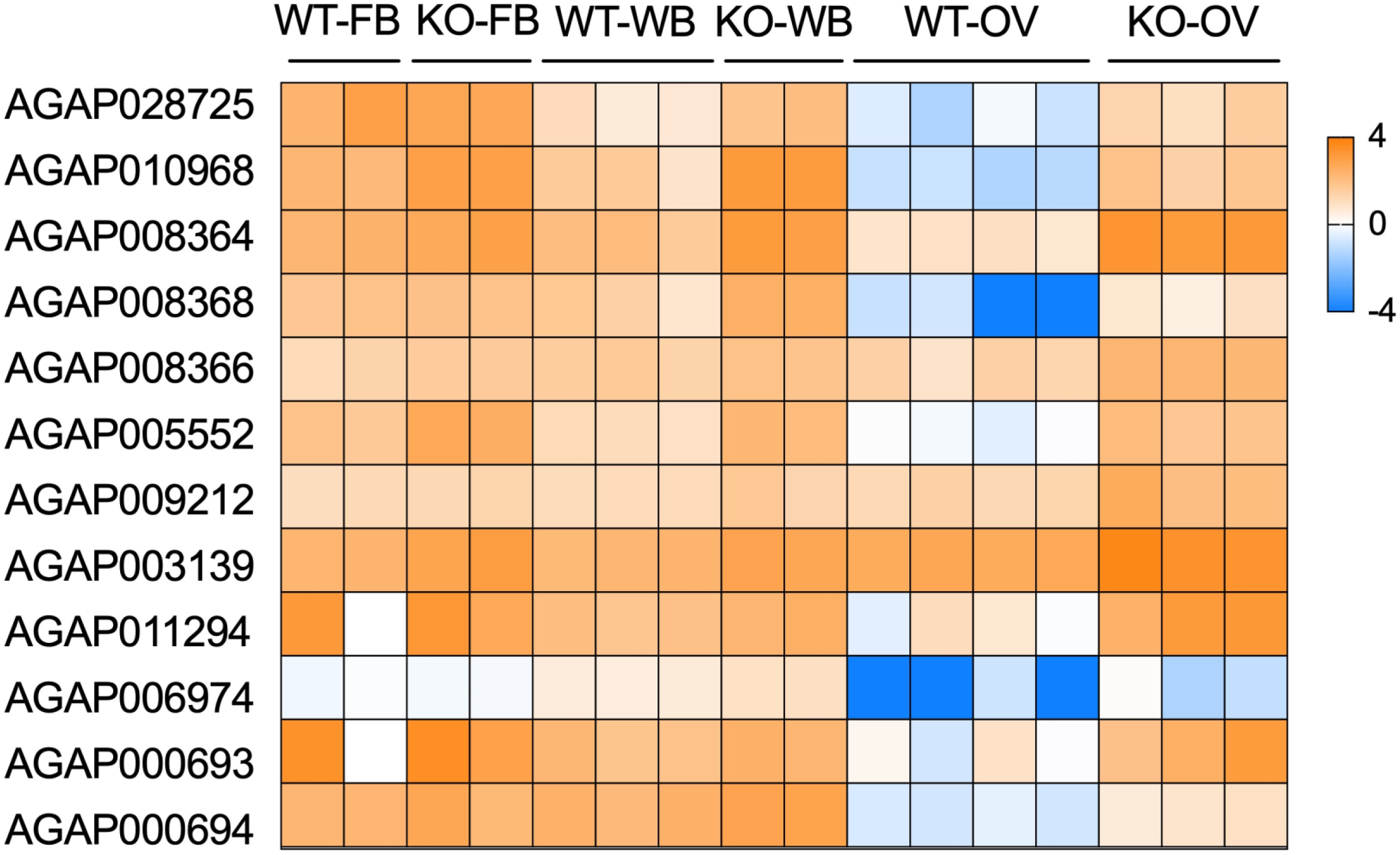
Heat map analysis of key differentially expressed genes related to immune pathways in the whole body (WB), fat body (FB) and ovary (OV) of *mosGILT^null^* mosquitoes.

In this study, we also found that *mosGILT^null^* mosquitoes have increased expression of three TEP proteins - TEP15, TEP14, and TEP2. TEP15 (AGAP008364) expression was increased in the ovary (fold change 230, p-value <0.0001) and whole body (fold change 12, p-value 0.0001) (Fig. 6). TEP14 (AGAP008368, fold change 10, p-value 0.0017) and TEP2 (AGAP008366, fold change 9, p-value 0.0001) expression was increased in the whole body. These *TEP* genes were upregulated significantly upon *P. falciparum* infection in the midgut of *A. gambiae* mosquitoes (Dong et al., 2006).

mosGILT disruption also increased the expression of another innate immune gene, peptidoglycan recognition protein, PGRPLD (AGAP005552) in the ovary (fold change 125, p-value <0.0001), whole body (fold change 8, p-value <0.0001) and fat body (fold change 7, p-value 0.0001). Peptidoglycan recognition proteins (PGRPs) are a family of immune regulators, conserved from insects to mammals. PGRPLD is important for anti-*Plasmodium* immunity and in *PGRPLD*-silenced mosquitoes, *Plasmodiu*m infection increased significantly (Gendrin et al., 2017). Additional immune-related genes, including 2 serpins, SRPN6 (AGAP009212) and SRPN9,(AGAP003139) was also overexpressed in the ovary (fold change 11, p-value 0.0002, fold change 8, p-value <0.0001 respectively) and whole body (fold change 3, p-value 0.0135, fold change 3, p-value 0.0002 respectively) and fat body (SRPN9, fold change 5, p-value 0.0007) (Fig. 6). SRPN6 is a component of the midgut epithelial immune-response system which is increased during *Plasmodium* infection and knockdown of this gene increased parasite numbers in *A. stephensi* (Abraham et al., 2005).

In addition, we also observed increase in defensin transcripts, an anti-microbial peptide (AGAP011294, DEF1). *Def1* expression was upregulated in the ovary (fold change 255, p-value 0.0001) and whole body (fold change 3, p-value 0.0029) of *mosGILT^null^* mosquitoes (Fig. 6). Transcripts of another anti-microbial peptide gene, *cercopin 1* (AGAP000693) was also increased in the ovary (fold change 300, p-value 0.0004) and whole body of the *mosGILT^null^* mosquitoes (fold change 2, p-value 0.0235), and *cercopin 3* (AGAP000694) expression was upregulated in the whole body (fold change 3, p-value 0.0016). Toll-like receptors (TLRs) are evolutionarily conserved pattern recognition receptor proteins with diverse biological functions. In *Drosophila melanogaster*, Toll plays an important role in pattern formation during embryogenesis and in immunity. In this study we observed that TLR9 expression in the *mosGILT^null^* mosquito was upregulated in the whole body (AGAP006974, fold change 2, p-value 0.0095) Though there is not much information on this gene in *A. gambiae*, in the silk moth, *Bombyx mori*, TLR9 is involved in innate immunity and functions as a pattern recognition receptor that binds lipopolysaccharide.

## Discussion

To further understand *Plasmodium* infection of the vector, and develop new mosquito and malaria control strategies, it is important to decipher the molecular mechanisms related to mosquito reproduction and immunity. Recently our lab uncovered a critical role for mosGILT in *A. gambiae* ovary development and *Plasmodium* infection (Yang et al., 2020). The *mosGILT^null^* mosquitoes displayed underdeveloped ovaries and reproductive defects including impaired production of 20E and Vg while showing an increasedTEP1-mediated immune response against *Plasmodium* ookinetes (Yang et al., 2020). However, there is a lack of knowledge regarding how mosGILT functions in *A. gambiae*, its interacting partners, and the implicated signaling pathways. Here we characterize the transcriptional response in *mosGILT^null^* mosquitoes, including the ovaries, whole body, and fat body, and delineate how mosGILT is regulating reproduction and immune responses.

GILT is a thioredoxin-related oxidoreductase which is the only known lysosomal thiol reductase (Balce et al., 2014). Mammalian GILT is related to the thioredoxin family of oxidoreductases, characterized by a CXXC motif in the active site and an enzymatic mechanism in which the pair of active site cysteine residues cooperate to reduce substrate disulfide bonds (Fomenko and Gladyshev, 2002; Hastings and Cresswell, 2011; Kongton et al., 2014; Schleicher et al., 2018). In *Anopheles* mosquitoes and other insect genera, the active motif is CXXS, where the second cysteine is replaced with serine (Fig. 7). This change in the active site of mosGILT hinders the thioredoxin reaction of the mixed intermolecular disulfide. The CXXS containing thioredoxins can interact with their substrate proteins in a more stable manner since there is a covalent complex formed between thioredoxins and their target. However, some studies report that at least some CXXS-containing forms of thioredoxin-fold proteins retain catalytic activity albeit at a lower level (Anelli et al., 2002; Norgaard and Winther, 2001). It is therefore possible that mosGILT has partial thioredoxin activity that is critical to the transcriptional response that leads to oogenesis and ovarian development.

**Figure 7:**
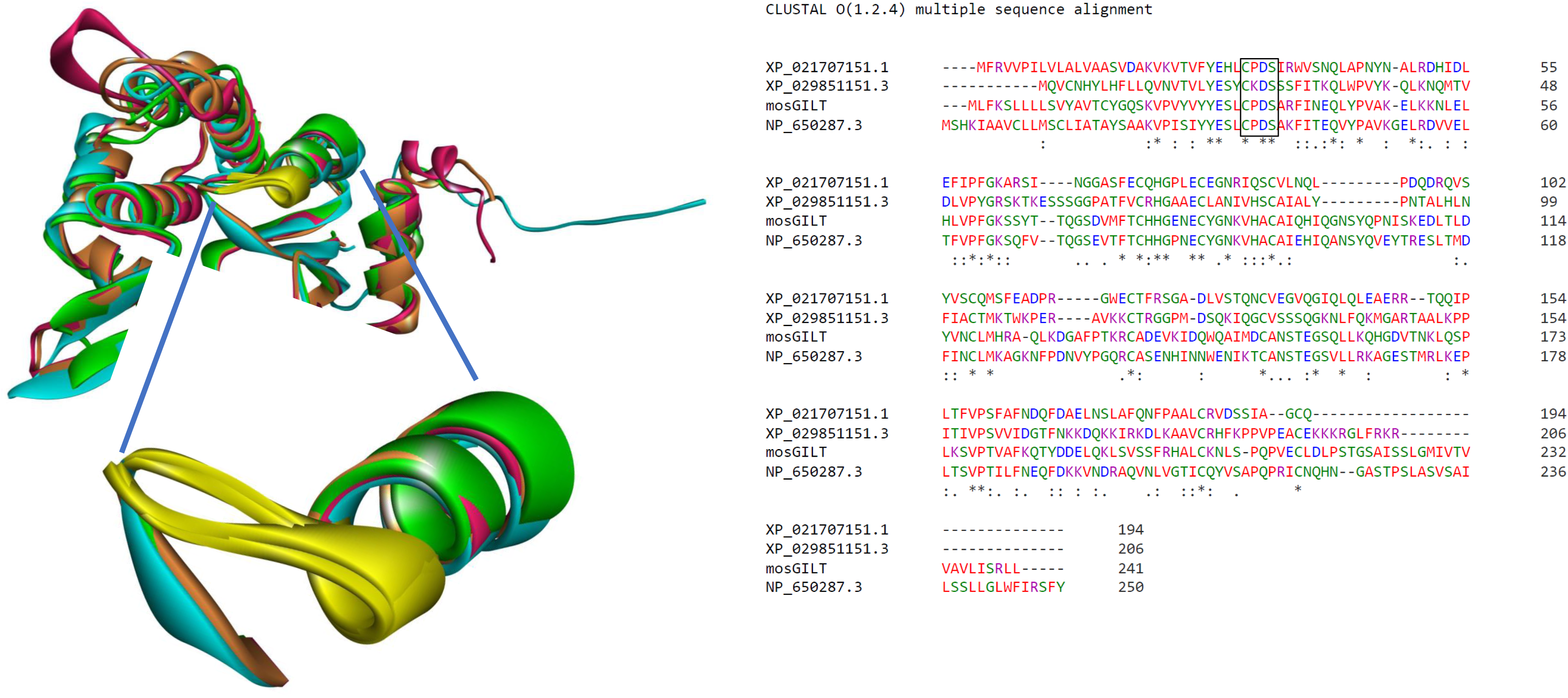
Structural and sequence analysis of mosGILT (A) The mosGILT structural model is created by Alpha Fold. Structural alignment of mosGILT homologs from *Anopheles*, *Aedes*, *Drosophila* and human GILT. The active site motif is shown in the yellow in the inset. (B) Sequence alignment of *Anopheles*, *Aedes*, *Drosophila* and *Ixodes* GILT. Protein sequence of mosGILT homologs from *Anopheles gambiae* (AGAP004551), *Aedes aegypti* (XP_021707151.1 GILT-like protein 1), *Drosophila melanogaster* (NP_650287.3 Gamma-interferon-inducible lysosomal thiol reductase 1), *Ixodes scapularis* (XP_029851151.3 GILT-like protein 1). Active site residues are shown in the inset box.

Vector reproduction intimately interacts with immune defense pathways, as these two processes are simultaneously initiated after a blood meal (Clayton et al., 2014; Reynolds et al., 2020; Upton et al., 2015). Identification of vector proteins involved in the cross-talk between reproduction and immunity could provide new targets for the development of disease-control strategies. Our study demonstrates the critical role of mosGILT in the reproduction of *A. gambiae*. Our study also shows that mosGILT influences complement activity and upregulates defense-related genes, which are associated with anti-*Plasmodium* responses. In addition, mosGILT regulates TEP1 activity by altering SPCLIP1 expression, which can influence *Plasmodium* survival in mosquitoes. *mosGILT^null^* mosquito refractoriness to microbes was also suggested by the increase in genes encoding antimicrobial peptides and innate immune molecules. *mosGILT* may therefore be a potential target for new malaria control strategies, such as gene drive -propelled transgenes targeting mosGILT, that would induce mosquito sterility and also reduce the overall malaria parasite burden.

*mosGILT* mutants have underdeveloped ovaries that produced low amounts of 20E, and were significantly less permissive to *P. falciparum* infection (Yang et al., 2020). This is somewhat similar to *Plasmodium* infection in *zpg* mutant mosquitoes, a gene that is essential for germ cell development (Tazuke et al., 2002; Werling et al., 2019). Though both mutant mosquitoes provide an association between 20E signaling and intensity of *Plasmodium* infection, it is not known if they act together or independently. Our studies now suggest that mosGILT may affect several genes that are associated with zpg. Since *zpg* and other germ cell development genes are considered a prime target for gene drive(Hammond et al., 2021; Kyrou et al., 2018; Terradas et al., 2022), our results suggest mosGILT may be more attractive target for gene drive of *A. gambiae*.

mosGILT plays an important role in reproduction and host defenses in *A. gambiae*. The elucidation of mosGILT as a regulator of *zpg* and involvement in cross-talk between reproduction and immunity could provide new targets for the development of disease-control strategies (Fig. 8). These findings can potentially lead to new preventive strategies for malaria based on *Anopheles* mosGILT and also potentially suggest a role for arthropod GILT-like proteins for other vector-borne diseases.

**Figure 8:**
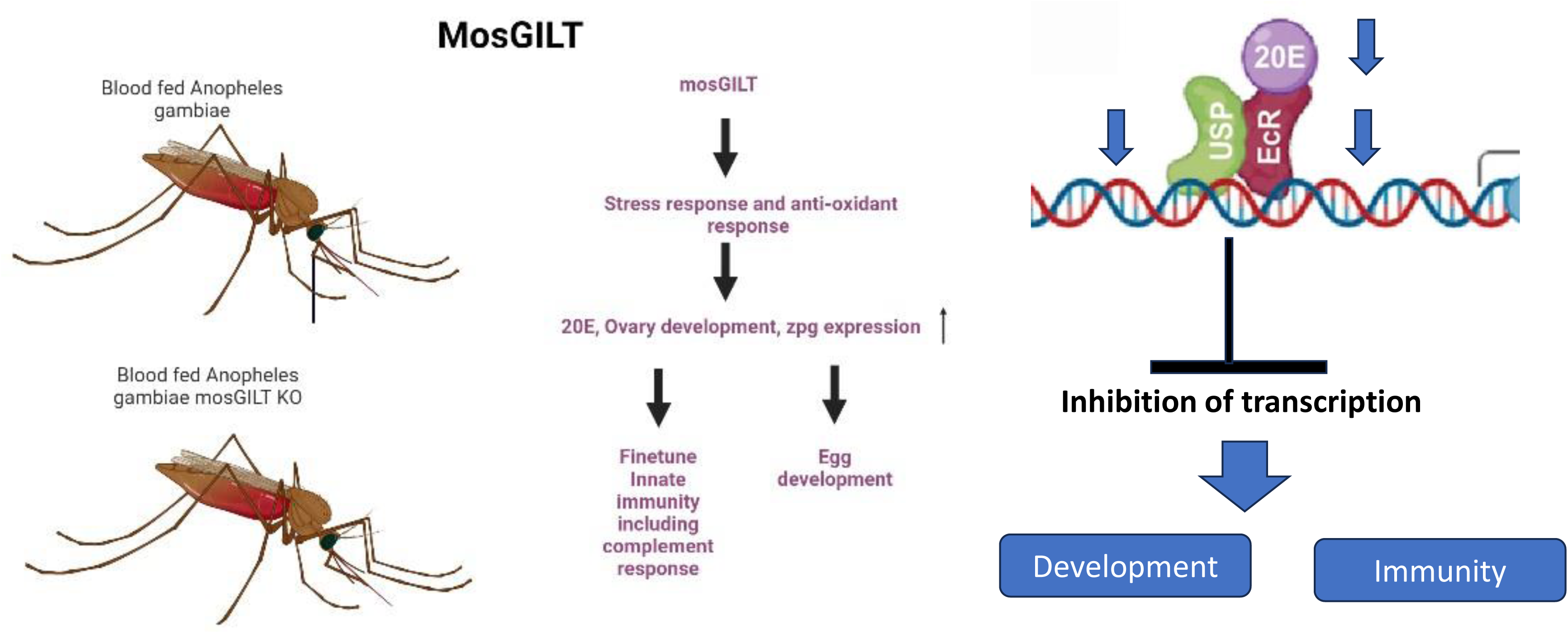
Diagrammatic representation of mosGILT’s role in immunity and development

## Materials and Methods

### Mosquito rearing

All the *A. gambiae* mosquito lines were derived from the G3 strain, including the docking line X1 (Volohonsky et al., 2015), *vasa2-Cas9* (Hammond et al., 2016), *mosGILT*-*gRNA_3_* (Yang et al., 2020), *mosGILT* mutant and sibling mosquitoes. They were raised at 27°C, 80% humidity, under a 12/12 h light/dark cycle and maintained with 10% sucrose. Swiss Webster mice (8-week-old females) were purchased from Charles River Laboratories.

### Generation of transgenic *mosGILT* mutant mosaic *Anopheles* mosquitoes

CRISPR/Cas9 mediated *A. gambiae mosGILT* somatic KO mutants were generated using the *A. gambiae* docking line X1 as described in Yang et al., 2020. In brief, the *mosGILT*-gRNA female mosquitoes were crossed with the homozygous male *Vasa Cas9* transgenic strain (*vasa2-Cas9*; with green/yellow fluorescence in the eyes; Hammond et al., 2016)(Dong et al., 2018). The progeny of this cross was screened at the larvae stage with both RFP (red fluorescence marker) and YFP (appeared as green fluorescence through GFP-channel under the fluorescence microscope) signals was *gRNA/Cas9* transheterozygous. Molecular screening of these transheterozygotes by PCR and sequencing was done as described to confirm the *mosGILT^null^* (Yang et al., 2020).

### Ethics statement

Mice used in the study were housed at the Yale Animal Resource Center of Yale University. Mice were handled in accordance with the Guide for the Care and Use of Laboratory Animals of the National Institutes of Health, USA. The Institutional Animal Care and Use Committee of Yale University approved the animal experimental protocol (protocol permit no. 2017–07941).

### RNA isolation

Total RNA was extracted from the whole body, ovaries, or fat bodies from mosquitoes. Trizol was added to the tissues and RNA was isolated using a using a combination method which utilizes TRIzol (Thermo Fisher Scientific) and RNeasy Mini Kit (QIAGEN) as described before (Untergasser, 2008).

### RNA-seq

RNA was submitted for library preparation using TruSeq (Illumina, San Diego, CA, USA) and sequenced using Illumina HiSeq 2500 by paired-end sequencing at the Yale Center for Genome Analysis (YCGA).

### Differential gene expression analysis

All the RNA-seq analyses including alignment, quantitation, normalization, and differential gene expression analyses were performed using Partek Genomics Flow software (St. Louis, MO, USA). Specifically, RNA-seq data were trimmed and aligned to the *Anopheles* PEST genome, version P4.14. with associated annotation file using STAR (v2.7.3a) (Dobin et al., 2013). The aligned reads were quantified using the Partek E/M algorithm (Xing et al., 2006) and the subsequent steps were performed on gene-level annotation followed by total count normalization. The gene-level data were normalized by dividing the gene counts by the total number of reads followed by the addition of a small offset (0.0001).

### Principal components analysis

Principal components analysis (PCA) was performed using default parameters for the determination of the component number, with all components contributing equally in Partek Flow. Volcano plot Hierarchal clustering was performed on the genes that were differentially expressed across the conditions (P < 0.05, fold change ≥ 2 for each comparison).

### Pathway enrichment analysis

Pathway enrichment was also conducted in Partek Flow and the functional annotation tool DAVID (https://david.ncifcrf.gov/tools.jsp). The selected genes expression heatmap was further plotted by ggplot2 (R sutdio) and Prism v8 (Graphpad). The selected immune pathways were further plotted on a bubble diagram by ggplot2 in R studio. The network analysis was performed in String database.

### PPI network and Cluster analysis

PPI network was generated by STRING online database and visualized by Cytoscape software. K means clustering with 10 clusters was selected to generate cluster with edges.

### Statistical analysis

All data analysis, graphing, and statistics were performed in Prism (v8.0, GraphPad Software) and R studio.

### Data availability

RNAseq data should be deposited to the public database (Sequence Read Archive (SRA) data,. The RNA-seq data are available in the Gene Expression Omnibus (GEO) repository at the National Center for Biotechnology Information under the accession number: GSE239600.

## Acknowledgements

We thank the Yale Center for Genome Analysis (YCGA) for conducting RNA sequencing. This work is supported in part by the Howard Hughes Medical Institute Emerging Pathogens Initiative and NIH grants (AI142708, and AI58615), and the Bloomberg Philanthropies. We thank the JHMRI insectary core facility for maintaining *A. gambiae* mosquitoes.

## Supplementary Figure Legends

**Supplementary Figure 1: Distribution of Raw counts**

**Supplementary Figure 2: Key statistics of Ovary samples**

**Supplementary Figure 3: Key statistics of whole body samples**

**Supplementary Figure 4: Key statistics of Fatbody samples**

**Supplementary Figure 5: Protein-Protein Interaction (PPI) network:** The interaction Network of DEGs in the Ovary of *mosGILT^null^* mosquitoes was created by STRING database and imported into Cytoscape. The k clustering method was used to identify clusters and key nodes in this network.

## Notes

### Competing Interest Statement

The authors have declared no competing interest.

