## Supplementary figures for "*Anopheles gambiae* mosGILT regulates innate immune genes and *zpg* expression"

### Supplementary Figure 1

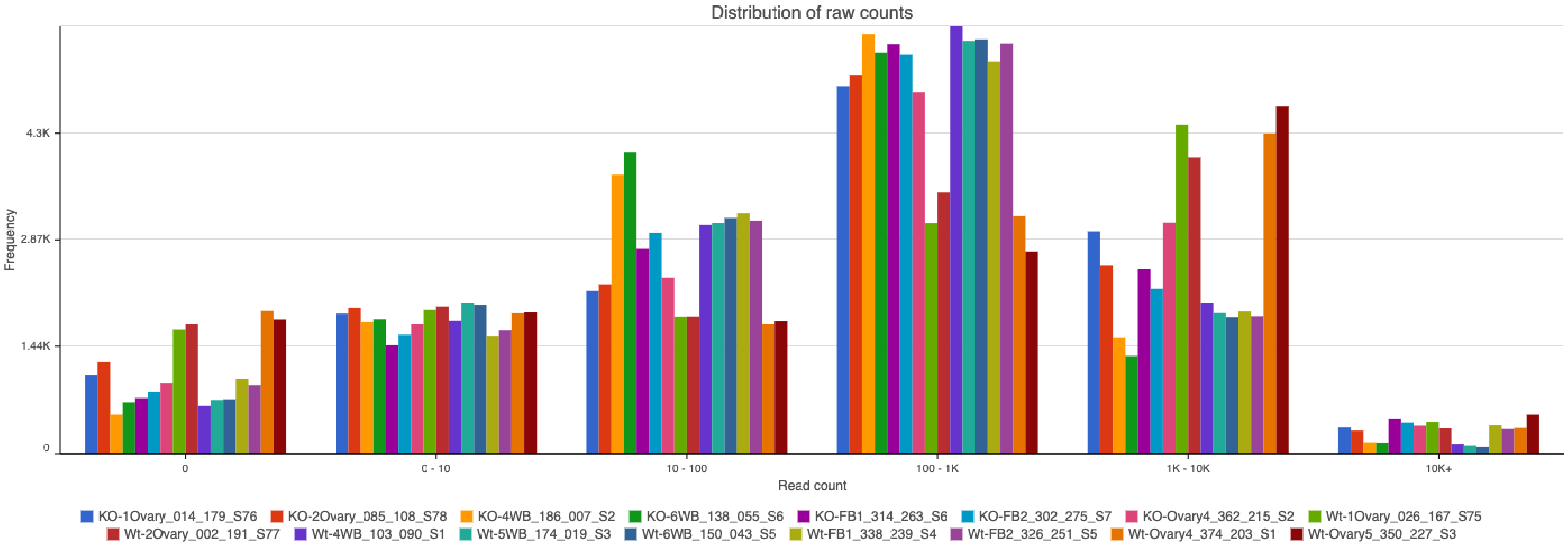

### Supplementary Figure 2: Key Statistics of Ovarian samples

**Summary of reads quantified to aga - gambiae**

Optional columns

| Sample name | Total reads | Fully within an exon | Partly within an exon | Fully within an intron | Fully intergenic | Incompatible paired-end | Compatible junctions | Total junctions | View |
|---|---|---|---|---|---|---|---|---|---|
| KO-1Ovary_014_179_S76 | 26,318,938.00 | 78.73% | 3.13% | 3.99% | 8.70% | 5.45% | 3,400,774.00 | 4,370,374.00 | |
| KO-2Ovary_085_108_S78 | 22,776,096.00 | 78.46% | 3.11% | 3.82% | 9.06% | 5.55% | 2,967,114.00 | 3,805,961.00 | |
| KO-Ovary4_362_215_S2 | 27,398,899.00 | 78.03% | 4.24% | 3.98% | 6.96% | 6.80% | 5,323,700.00 | 6,925,909.00 | |
| Wt-1Ovary_026_167_S75 | 28,468,342.00 | 82.18% | 3.05% | 2.39% | 6.89% | 5.49% | 3,601,721.00 | 4,561,748.00 | |
| Wt-2Ovary_002_191_S77 | 23,381,670.00 | 81.93% | 3.14% | 2.38% | 7.00% | 5.55% | 2,876,711.00 | 3,661,624.00 | |
| Wt-Ovary4_374_203_S1 | 25,073,627.00 | 81.30% | 4.52% | 2.09% | 5.93% | 6.16% | 4,222,895.00 | 5,357,611.00 | |
| Wt-Ovary5_350_227_S3 | 32,241,676.00 | 81.89% | 4.42% | 1.83% | 5.58% | 6.28% | 5,805,263.00 | 7,427,826.00 | |
| Average | 26,522,749.71 | 80.42% | 3.69% | 2.89% | 7.08% | 5.92% | 4,028,311.14 | 5,158,721.86 | |

Rows per page 25 (1 of 1) Download

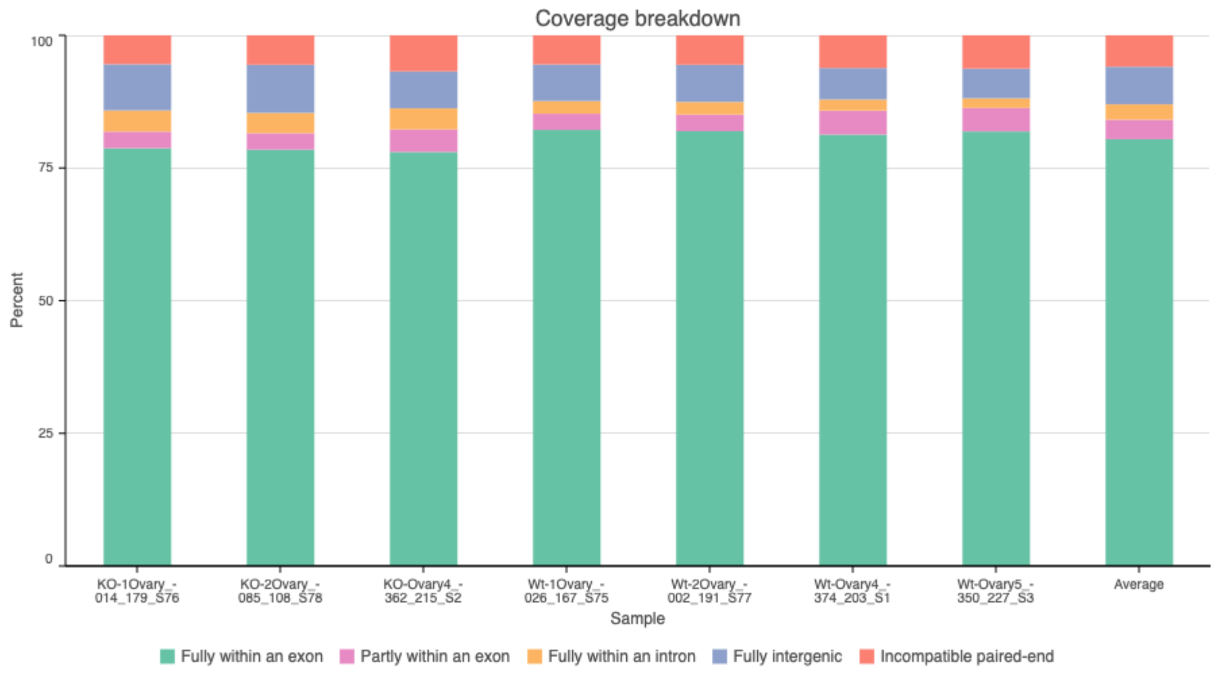


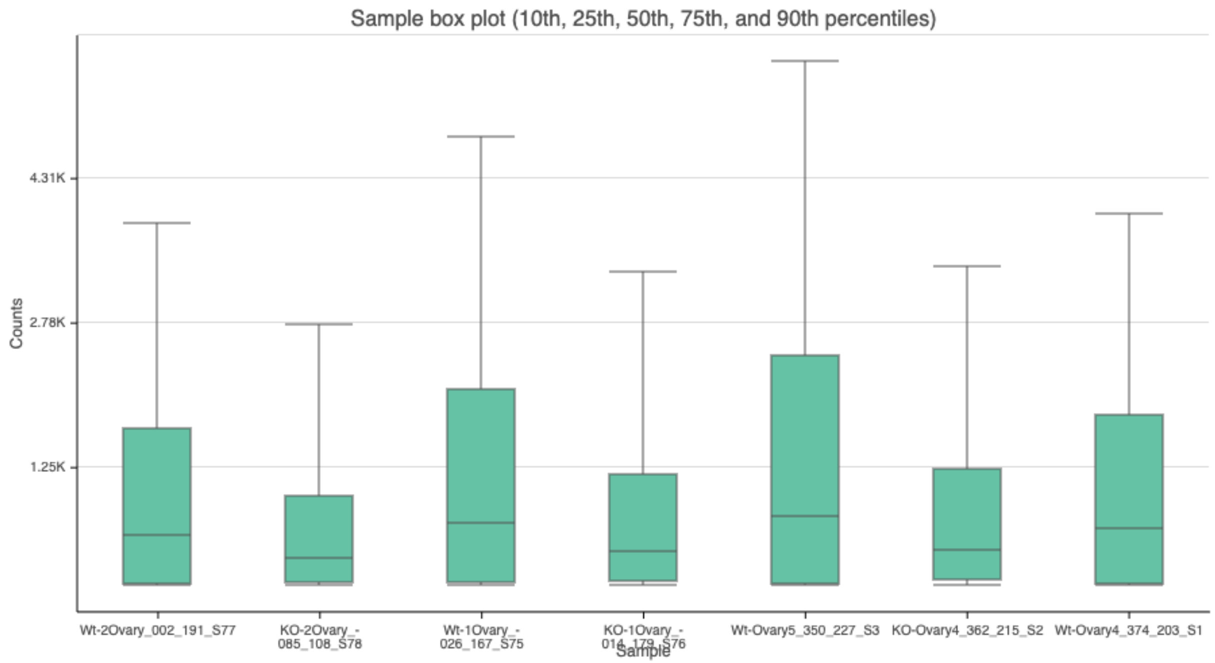

### Supplementary Figure 3: Key statistics of Whole body

Optional columns

| Sample name | Total reads | Fully within a feature | Partly within a feature | Not in a feature | Incompatible paired-end | Compatible junctions | Total junctions | View |
|---|---|---|---|---|---|---|---|---|
| KO-4WB_186_007_S2 | 10,712,966.00 | 99.74% | 0.10% | 0.00% | 0.16% | 2,252,491.00 | 2,280,500.00 | |
| KO-6WB_138_055_S6 | 10,799,903.00 | 99.76% | 0.10% | 0.00% | 0.14% | 1,996,089.00 | 2,021,807.00 | |
| Wt-4WB_103_090_S1 | 12,371,793.00 | 99.77% | 0.09% | 0.00% | 0.14% | 2,279,888.00 | 2,308,628.00 | |
| Wt-5WB_174_019_S3 | 9,958,040.00 | 99.72% | 0.11% | 0.00% | 0.17% | 2,102,655.00 | 2,130,788.00 | |
| Wt-6WB_150_043_S5 | 9,436,270.00 | 99.72% | 0.11% | 0.00% | 0.17% | 1,801,246.00 | 1,827,209.00 | |
| Average | 10,655,794.40 | 99.74% | 0.10% | 0.00% | 0.15% | 2,086,473.80 | 2,113,786.40 | |

Rows per page 25 (1 of 1) Download

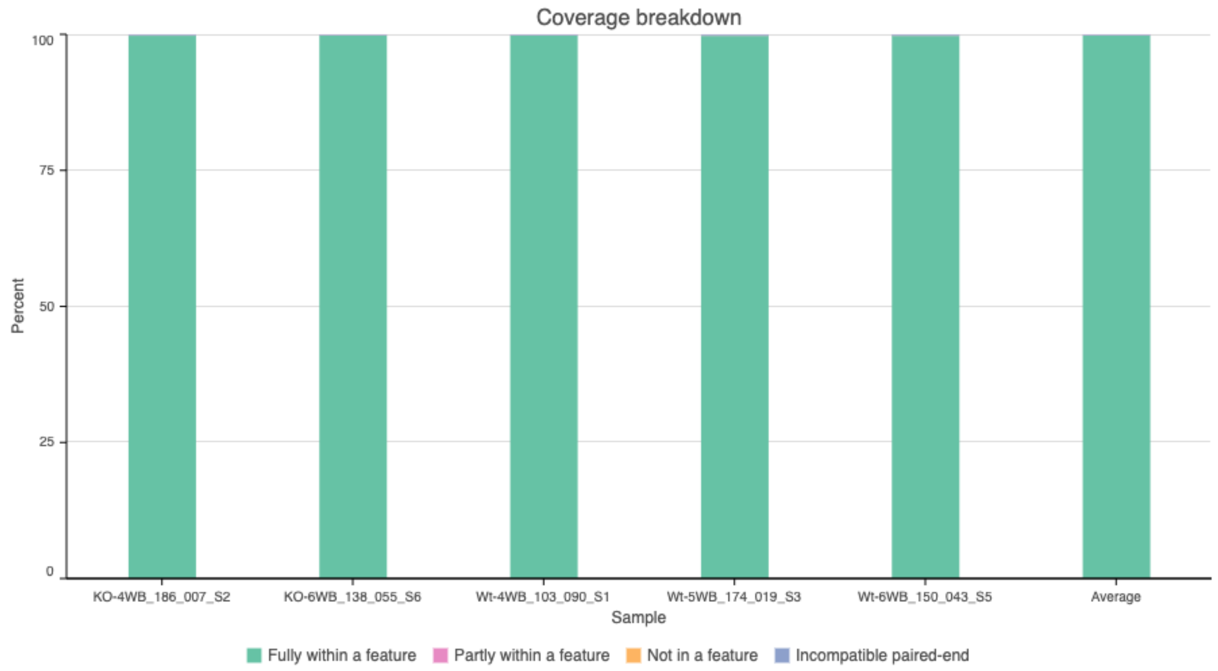


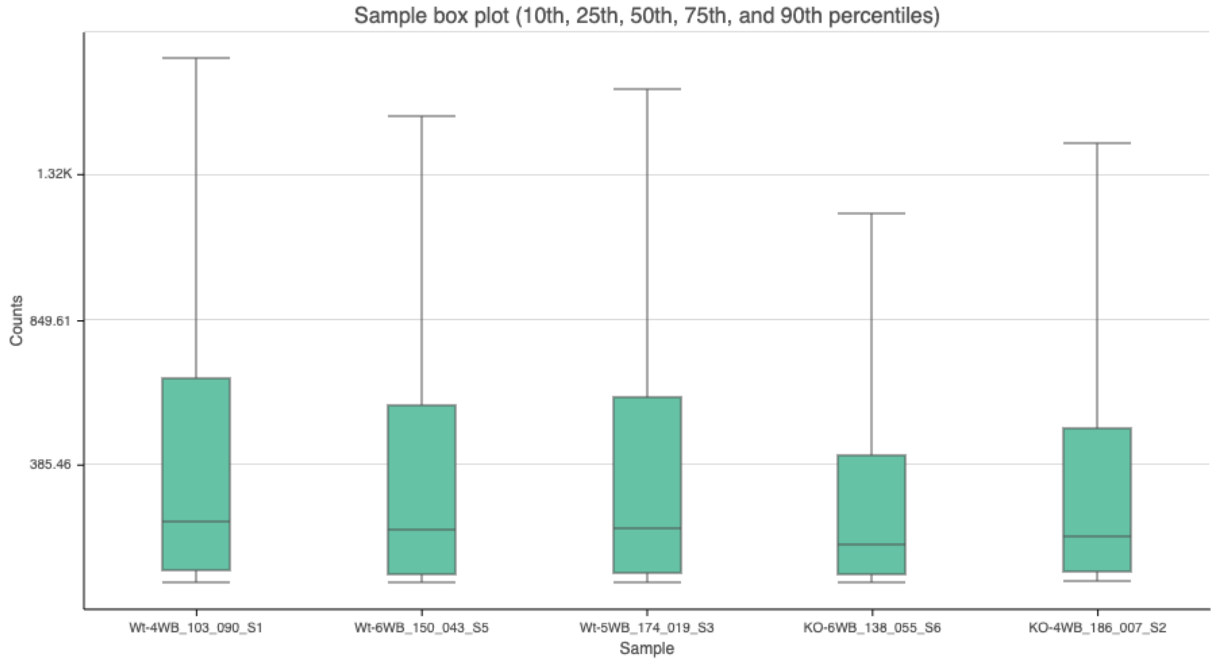

### Supplementary Figure 4: Key statistics of Fatbody samples

Optional columns

| Sample name | Total reads | Fully within a feature | Partly within a feature | Not in a feature | Incompatible paired-end | Compatible junctions | Total junctions | View |
|---|---|---|---|---|---|---|---|---|
| KO-FB1_314_263_S6 | 29,746,900.00 | 82.95% | 3.68% | 7.27% | 6.10% | 5,293,676.00 | 6,685,353.00 | |
| KO-FB2_302_275_S7 | 30,136,606.00 | 84.56% | 3.54% | 5.38% | 6.53% | 5,591,036.00 | 7,102,196.00 | |
| Wt-FB1_338_239_S4 | 28,560,821.00 | 83.39% | 3.66% | 6.50% | 6.44% | 4,362,502.00 | 5,486,092.00 | |
| Wt-FB2_326_251_S5 | 26,714,071.00 | 83.84% | 3.68% | 5.79% | 6.69% | 4,050,396.00 | 5,172,691.00 | |
| Average | 28,789,599.50 | 83.69% | 3.64% | 6.24% | 6.43% | 4,824,402.50 | 6,111,583.00 | |

Rows per page 25 (1 of 1) Download

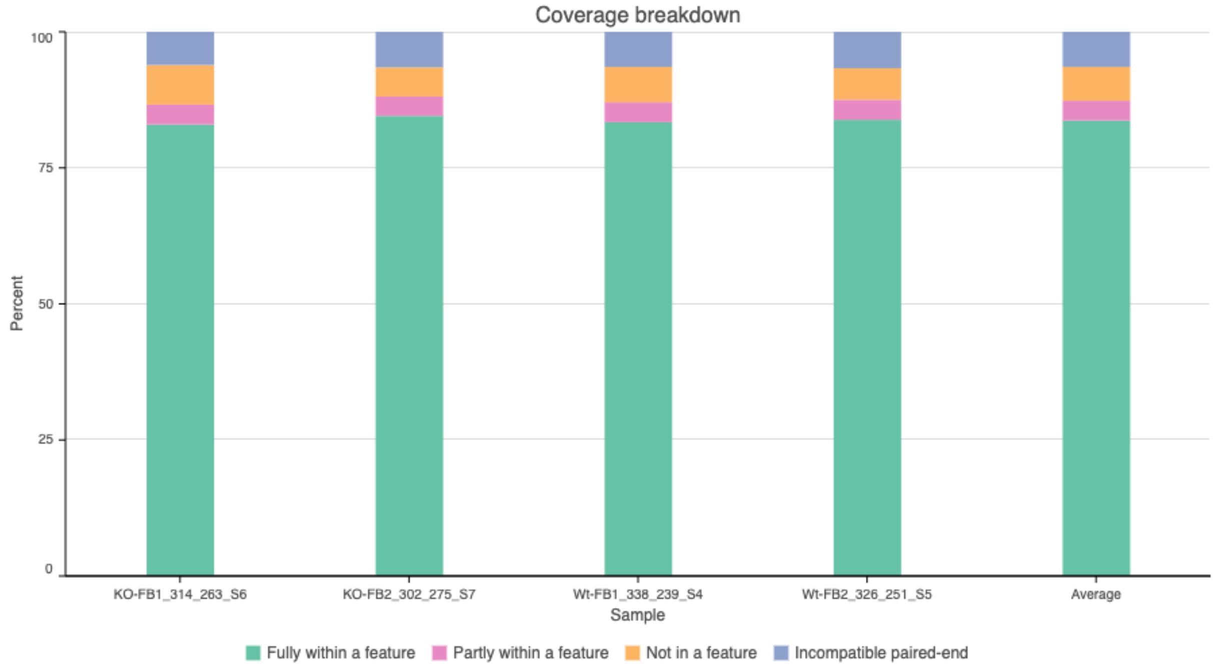


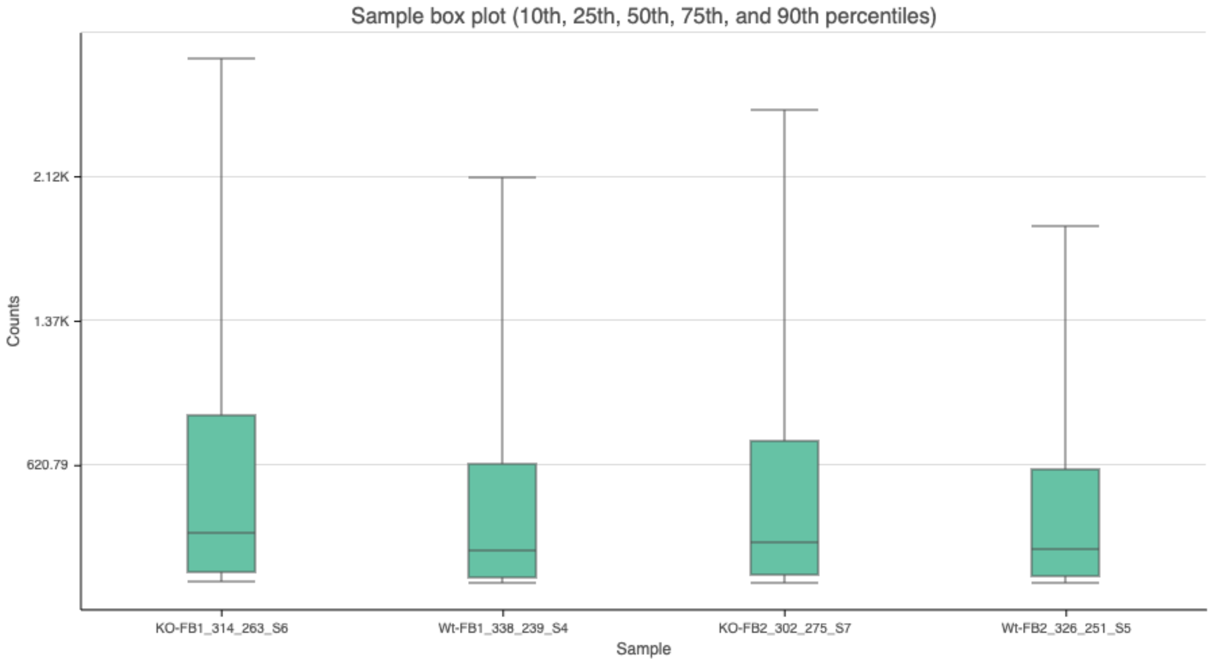

### Supplementary Figure 5

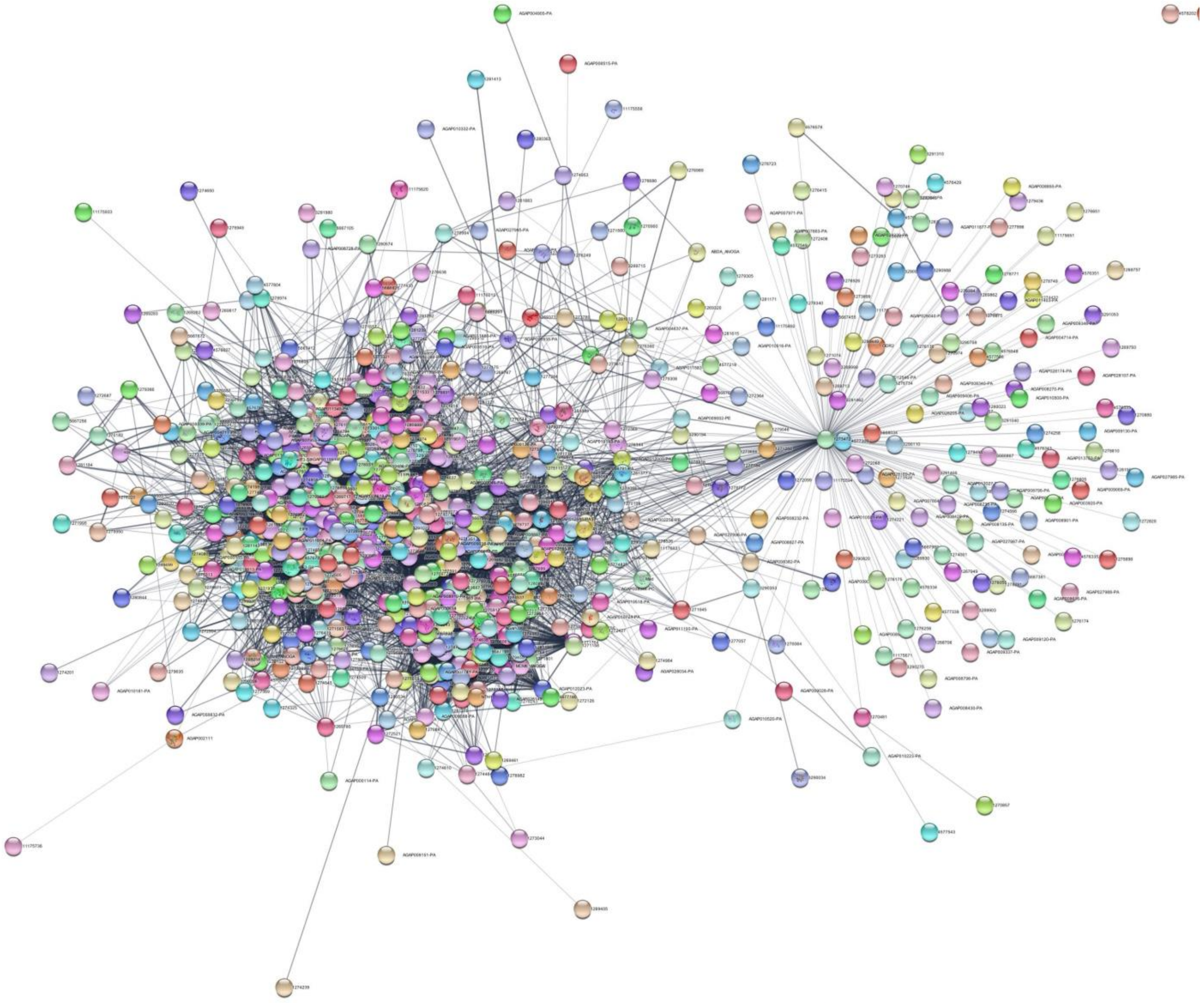

**Supplementary Figure 5:** **Protein-Protein Interaction (PPI) network:** The interaction Network on DEGs in the Ovary of *mosGILT-KO* mosquitoes was created by STRING database and imported into Cytoscape. The k clustering method was used to identify clusters and key nodes in this network.

### Supplementary Table 1: Separate as excel sheet
